# Phosphorylation and Lamin B receptor modulate the accumulation of PERIOD protein foci

**DOI:** 10.1101/2023.01.16.524233

**Authors:** Mengna Li, Shujing Li, Luoying Zhang

## Abstract

Circadian clock drives the 24h rhythm in our behavior and physiology. Previously, we have shown that inner nuclear membrane protein Lamin B receptor (LBR) regulates circadian rhythm in human cells and *Drosophila*, while the underlying mechanism is unclear (Lin et al., 2014). A very recent study reported that the circadian clock protein PERIOD is organized into discrete foci at the nuclear envelope in fly circadian neurons, which is believed to be important for controlling the subcellular localization of clock genes. Loss of LBR leads to disruption of these foci, but how they are regulated is yet unknown. Here we found that LBR likely facilitates PER foci accumulation by destabilizing the catalytic subunit of protein phosphatase 2A, MICROTUBULE STAR (MTS). MTS is known to dephosphorylate PER and hampers the accumulation of PER foci. On the other hand, the circadian kinase DOUBLETIME (DBT) which phosphorylates PER enhances the accumulation of the foci. These foci are likely phase-separated condensates, the formation of which mediated by intrinsically disordered region in PER. Taken together, here we demonstrate a key role for phosphorylation in promoting the accumulation of PER foci, while LBR modulates this process by impinging on the circadian phosphatase MTS.

## Introduction

Most if not all organisms on the earth display circadian rhythms, manifested in various aspects of physiology and behavior. These rhythms are driven by endogenous circadian clocks, which at the molecular level consist of a series of transcriptional/translational feedback loops (TTFLs) operated by a number of core clock genes (Li & Zhang, 2015). The molecular clock orchestrates rhythmic transcription of a significant portion of the genome, ranging from ∼1-60%, depending on the species and organ/tissue type. Although the regulatory mechanism underlying the molecular clock itself is fairly well-characterized, how it cyclically modulates global transcription remains largely unclear. Previous studies have reported the presence of circadian long-range interactions within the genome, and that chromosomes are rhythmically recruited to the nuclear lamina which leads to transcription attenuation (Aguilar-Arnal et al., 2013; Zhao et al., 2015). Our work have also demonstrated that the inner nuclear membrane protein MAN1 regulates the transcription of core clock genes in mice, flies, and human cells (Bu, He, Song, & Zhang, 2019; Lin et al., 2014). However, the mechanism by which the nuclear lamina regulates the spatial-temporal organization of the chromosome and rhythms in global gene transcription is unknown.

Very recently, Xiao et al discovered that core clock protein PERIOD (PER) forms discrete foci near the nuclear lamina in fly circadian neurons (Xiao, Yuan, Jimenez, Soni, & Yadlapalli, 2021). These foci are circadian controlled and may play a role in positioning core clock genes near the nuclear periphery where their transcription is repressed. Loss of lamin B receptor (LBR), an inner nuclear membrane protein, leads to disruption of PER foci formation and the nuclear periphery localization of the *per* gene. Our previous work has also reported that LBR regulates circadian rhythm in flies and human cells, but the mechanism has not been characterized (Lin et al., 2014). These findings indicate that understanding how LBR regulates the formation of PER foci may reveal some insights regarding how nuclear lamina contributes to the spatial-temporal organization of the chromosomes.

Here we found that LBR binds to the catalytic subunit of protein phosphatase 2A (PP2A), MICROTUBULE STAR (MTS). PP2A is known to dephosphorylate and stabilize PER protein, whereas here we demonstrate that LBR destabilizes MTS specifically in the nucleus (Sathyanarayanan, Zheng, Xiao, & Sehgal, 2004). MTS acts to hamper the accumulation of PER foci both *in vivo* and in cultured cells, whereas inhibiting MTS facilitates PER foci accumulation. These findings lead us to suspect that phosphorylation promotes the accumulation of PER foci, and indeed over-expressing DOUBLETIME (DBT), a circadian kinase known to phosphorylate PER, enhances PER foci accumulation (J. L. Price, Blau, J., Rothenfluh, A., Abodeely, M., Kloss, B., and Young, M.W., 1998). Further characterizations strongly suggest that these PER foci are phase-separated condensates mediated by intrinsic disordered region (IDR) in PER. Taken together, these results unveil a mechanism centering on phosphorylation which facilitates PER foci accumulation, while LBR participates in this process by influencing the stability of MTS.

## Results

### LBR binds to and destabilizes MTS

To investigate how LBR regulates PER foci formation, we first tested for direct physical interaction between LBR and PER. We co-expressed HA-tagged LBR and PER in fly S2 cells and performed immunoprecipitation using HA antibody but were not able to detect PER in the precipitates (Figure 1-figure supplement 1). By searching the literature, we found that PP2A, which is known to dephosphorylate PER and regulate circadian rhythm in flies, interacts with various components of the inner nuclear membrane including lamin, the binding partner of LBR (Mehsen et al., 2018; Nikolakaki, Mylonis, & Giannakouros, 2017; Van Berlo et al., 2005). Therefore, we suspected that instead of directly regulating PER, LBR may exert effects on PER via PP2A. Consistent with this hypothesis, we observed in S2 cells that LBR co-precipitates with MTS, the catalytic subunit of PP2A (Figure 1A). In flies with *lbr* knocked-down in all clock cells using a *timeless*(*tim*)GAL4 driver (Emery, So, Kaneko, Hall, & Rosbash, 1998), MTS protein level is significantly increased at the beginning of the day (Figure 1B-D). This elevation in protein level appears to be caused by alteration at the post-transcriptional level, as the mRNA level of *mts* is not significantly altered. We further probed the mechanism by which LBR regulates MTS by treating S2 cells with cycloheximide which inhibits protein synthesis, so we can observe the influences of LBR on MTS stability. We found that knocking down *lbr* delays MTS degradation in the nucleus but not the cytoplasm (Figure 1E-G). On the other hand, over-expressing *lbr* hastens MTS degradation in the nucleus but not the cytoplasm (Figure 1H-J). Taken together, this series of results indicate that LBR acts to destabilize MTS in the nucleus, and this effect appears to be most prominent early in the day.

**Figure 1.**
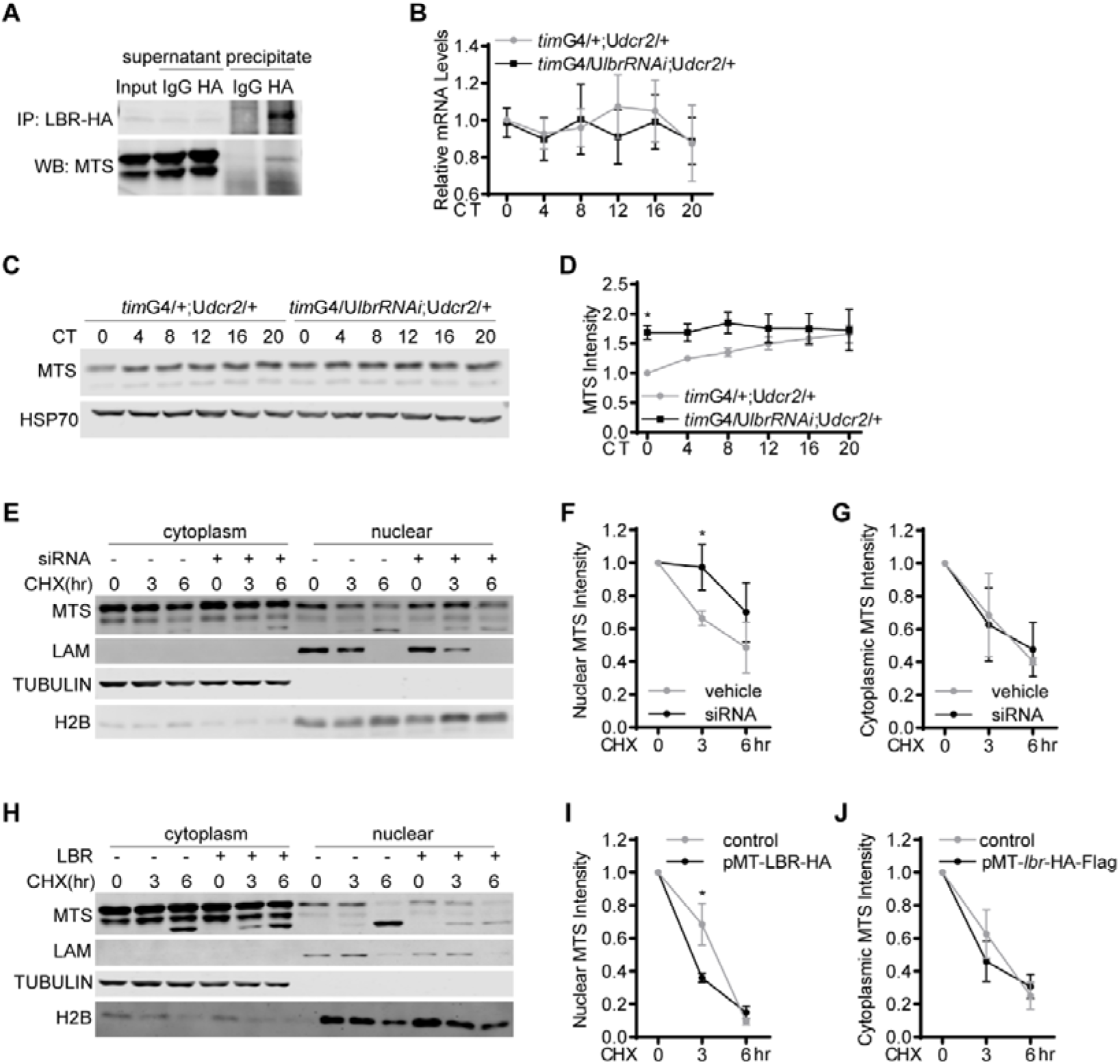
LBR binds to and destabilizes MTS. **(A)** Representative Western blots (WB) of protein extracts prepared from S2 cells transfected with pMT-*lbr*-Flag-HA and immunoprecipitates as well as supernatants. LBR was immunoprecipitated (IP) with HA antibody, and rabbit IgG was used as control. MTS was detected by Western blotting using MTS antibody. **(B)** Plot of relative mRNA abundance determined by qRT-PCR for *mts* from whole head extracts of *tim*GAL4/+;UAS*dcr2*/+ and *tim*GAL4/UAS*lbrRNAi*;UAS*dcr2*/+ flies collected on the first day of constant darkness (DD1). *dcr2* was co-expressed to enhance the efficiency of RNAi. For each time series, the value of control group at CT0 was set to 1. **(C)** Representative Western blots of protein extracts prepared whole heads of *tim*G4/+; U*dcr2*/+ and *tim*G4/U*lbrRNAi*;U*dcr2*/+ flies collected on DD1. HSP70 was used as a loading control. **(D)** Quantification of MTS level in (C), which was normalized to that of HSP70. For each group, the value of the control group at CT0 was set to 1. **(E)** Representative Western blots of cytoplasmic and nuclear extracts prepared from S2 cells transfected with *lbr* siRNA or control cells. The cells were treated with cycloheximide (CHX, 10μg/ml), and harvested at the indicated time points. (**F and G**) Quantification of MTS level in nuclear (**F**) and cytoplasmic (**G**) fraction in (E). (**H**) Representative Western blots of cytoplasmic and nuclear extracts prepared from S2 cells transfected with pMT-*lbr*-Flag-HA or control cells. The cells were treated with cycloheximide (10μg/ml) and harvested at the indicated time points. (**I and J**) Quantification of MTS level in nuclear (**F**) and cytoplasmic (**G**) fraction in (H). n=3. Error bars represent standard error of the mean (SEM). Two-way ANOVA, Sidak’s multiple comparison test was used for (B and D). Student’s t-test was used for (F, G, I and J). \**P* <0.05, \*\**P* <0.01, \*\*\**P* <0.001. G, GAL. U, UAS. This figure includes the following source data: **Figure 1-source data 1**. The original Western blots in ***Figure 1A, C, E and H*. Figure 1-source data 2**. The raw data of mRNA and protein levels as well as statistical analyses in ***Figure 1B, D, F, G, I and J***.

### MTS reduces PER foci accumulation

Since LBR appears to target MTS, we tested whether MTS regulates PER foci formation. Here we focused on PER foci by monitoring PER-EGFP in the small ventral lateral neurons (s-LNvs) which are considered to be the master pacemaker neurons signal (Chen, Werdann, & Zhang, 2018; Grima, Chelot, Xia, & Rouyer, 2004; Renn, Park, Rosbash, Hall, & Taghert, 1999; Stoleru, Peng, Agosto, & Rosbash, 2004). We over-expressed *mts* in these cells using a *pigment dispersing factor* (*Pdf*)GAL4 and assessed the size, intensity and number of PER foci at Circadian Time 0 (CT 0, defined as the time of subjective lights on) on the first day of constant darkness, as it has been reported that the foci show most prominent accumulation at this time point (Hannus, Feiguin, Heisenberg, & Eaton, 2002; Renn et al., 1999; Xiao et al., 2021). We found that over-expressing *mts* significantly reduced the size and number of the foci (Figure 2A and B). We also tried to inhibit MTS function by expressing a dominant negative form of *mts* (dn*mts*), but in these flies PER-EGFP signal is barely detectable due to reduction on PER stability as previously reported (Figure 2-figure supplement 1)(Hannus et al., 2002; Sathyanarayanan et al., 2004). To resolve this issue, we treated fly brains with phosphatase inhibitor calyculin A (Cal A) to assess the effects of acute PP2A inhibition on PER foci (Sathyanarayanan et al., 2004). This treatment significantly increased the size of PER foci with a trend of increase of foci number (Figure 2C and D), opposite to that of *mts* over-expression.

**Figure 2.**
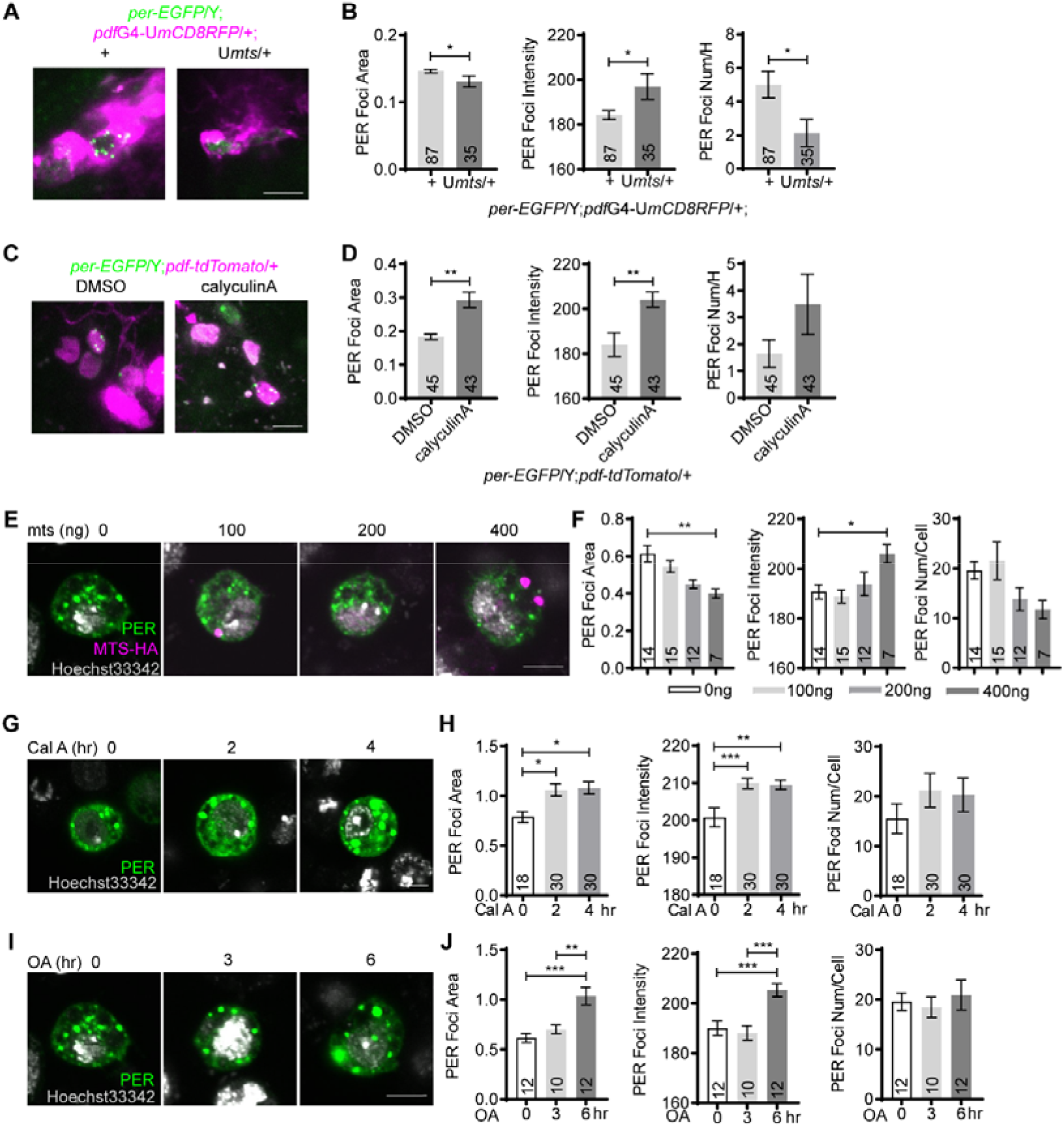
MTS reduces PER foci accumulation. **(A)** Representative confocal images of PER foci in the s-LNvs of *per*-EGFP/Y;*pdf*GAL4*-*UASmCD8RFP*/+* flies over-expressing *mts* and controls dissected at CT0 on DD1. Green, PER-EGFP; magenta, mCD8RFP. **(B)** Quantification of PER foci area, intensity and number per hemisphere (num/H) in (**A**). **(C)** Representative confocal images of PER foci in the s-LNvs of *per*-EGFP/Y;*pdf-*tdTomato/+ flies dissected at CT0 on DD1. The brains were incubated in PBS solution containing DMSO or 30nM calyculinA for 1hr. Green, PER-EGFP; magenta, tdTOMATO. **(D)** Quantification of PER foci area, intensity and number per hemisphere in (C). **(E)** Representative confocal images of PER foci in S2 cells co-transfected with 100ng pAc-*per* and indicated dosage of pMT-*mts*-HA for 36hr. The cells were then immune-stained with hoechst33342 (gray), PER antibody (green) and HA antibody (magenta). **(F)** Quantification of the PER foci area, intensity, number per cell in (E). **(G)** Representative confocal images of PER foci in S2 cells transfected with 100ng *per* for 30hr and then treated with calyculinA (Cal A, 30nM) for indicated time periods. The cells were subsequently immune-stained with hoechst33342 (gray) and PER antibody (green). **(H)** Quantification of PER foci area, intensity, number per cell in (G). **(I)** Representative confocal images of PER foci in S2 cells transfected with 100ng *per* for 30hr and treated with okadaic acid (OA, 5nM) for indicated time periods. The cells were then immunostained with hoechst33342 (gray) and PER antibody (green). **(J)** Quantification of PER foci area, intensity, number per cell in (I). Scale bar, 5μm. n number is indicated on the bars. For (B and D), n refers to the number of hemispheres. For (F, H, and J), n refers to the number of cells. Error bars represent SEM. Student’s t-test was used in (B and D). One-way ANOVA, Tukey’s multiple comparison test was used in (F, H and J). *\*P* <0.05, *\*\*P* <0.01, \*\*\**P* <0.001. G, GAL. U, UAS. This figure includes the following source data: **Figure 2-source data**. The raw data of PER foci area, intensity and number, as well as statistical analyses in ***Figure 2 B, D, F, H and J***.

We further validated these findings in S2 cells. A previous study has reported that PER accumulates in discrete foci in the cytoplasm of S2 cells (Meyer, 2006), and here we also observe PER foci both by immunostaining and live imaging (Figure 2-figure supplement 2A). To verify that these foci are similar to those *in vivo* and are not merely an artifact of over-expression, we treated cells with 1,6-hexanediol which disrupts weak hydrophobic interactions and found this leads to disassembly of the foci (Figure 2-figure supplement 2B)(Shin & Brangwynne, 2017). These results indicate that the PER foci in S2 cells display liquid-like properties, similar to what has been reported for the foci *in vivo* (Xiao et al., 2021).

Over-expressing *mts* in S2 cells significantly reduces the size of the foci in a dose-dependent manner, while foci number also exhibits a trend of reduction (Figure 2E and F). In contrast, Cal A treatment significantly increases the size and intensity of the foci, with a tendency of increase for foci number (Figure 2G and H). In addition, we treated cells with okadaic acid which inhibits PP2A with high affinity (Takai, Ohno, Yasumoto, & Mieskes, 1992), and this also leads to enhanced foci size and intensity (Figure 2I and J). Taken together, these findings indicate that MTS functions to impede the accumulation of PER foci.

### DBT enhances PER foci accumulation

The influences of MTS on PER foci lead us to suspect that phosphorylation promotes the accumulation of PER foci, therefore we tested whether DBT, the major kinase that phosphorylates PER, modulates PER foci accumulation (J. L. Price, Blau, J., Rothenfluh, A., Abodeely, M., Kloss, B., and Young, M.W., 1998). Over-expressing *dbt* in the s-LNvs significantly enlarges the size of PER foci (Figure 3A and B). Next, we over-expressed *dbt* in S2 cells and found this results in significant enhancement of foci size and intensity (Figure 3C and D). Interestingly, we also observed a striking pattern of PER and DBT co-localization in the foci.

**Figure 3.**
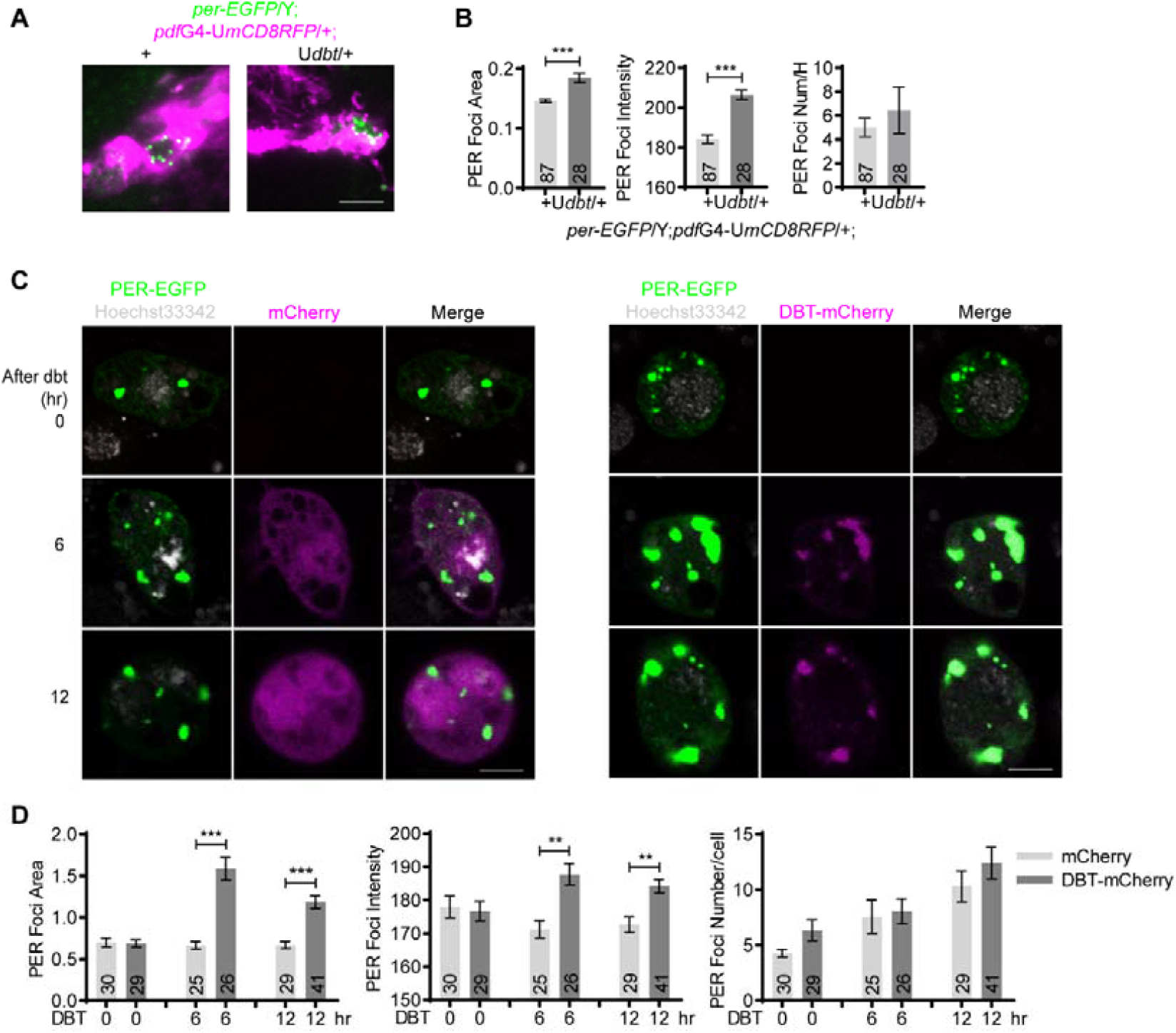
DBT increases PER foci size. **(A)** Representative confocal images of PER foci in the s-LNvs of *per-*EGFP/Y;*pdf*GAL4*-*UASmCD8RFP*/+* flies over-expressing *dbt* and control flies at CT0 on DD1. Green, PER-EGFP; magenta, mCD8RFP. **(B)** Quantification of PER foci area, intensity and number per hemisphere in (**A**). **(C)** Representative confocal images of PER foci in S2 cells transfected with 100ng pAc-*per*-EGFP and pMT-mCherry or pMT-*dbt-*mCherry for 30hr. Then 500μM CuSO4 was added to induce mCherry or *dbt-*mCherry expression. PER foci were live imaged at indicated time post induction. **(D)** Quantification of PER foci area, intensity and number of per cell in (**C**). Scale bar, 5μm. n refers to the number of cells and is indicated on the bars. Error bars represent SEM. Student’s t-test was used for (B). One-way ANOVA, Tukey’s multiple comparison test was used for (D). \*\**P* <0.01, \*\*\**P* <0.001. This figure includes the following source data: **Figure 3-source data**. The raw data of PER foci area, intensity and number, as well as statistical analyses in ***Figure 3B and D***.

### PER IDR form phase-separated condensates

We further investigated the liquid-like properties of PER foci by measuring the rate of fluorescence recovery after photobleaching (FRAP). We performed FRAP experiments on PER-EGFP foci in S2 cells. After photobleaching, fluorescence partially recovered on scale of seconds (Figure 4A and Figure 4-figure supplement 1), which indicates that they are dynamic. Since PER bears multiple IDRs as shown in past studies and IDRs can facilitate condensate formation, we tested whether these PER foci are phase-separated condensates (Fu et al., 2016; Shin & Brangwynne, 2017; Xiao et al., 2021). We selected the longest IDR (1-193), fused it to EGFP and expressed and purified it in a bacterial system (Figure 4-figure supplement 2A and B). EGFP and IDR-EGFP were added to 10% polyethelene glycol-6000 (PEG-6000) solution which increases macromolecular crowding and promotes phase-separation. IDR-EGFP rendered the solution opaque while EGFP solution remained clear (Figure 4B). We examined the solutions and observed EGFP-positive micron-sized droplets (Figure 4C). Live imaging demonstrates that the droplets are highly dynamic and two droplets can fuse into one (Figure 4-figure supplement 3).

**Figure 4.**
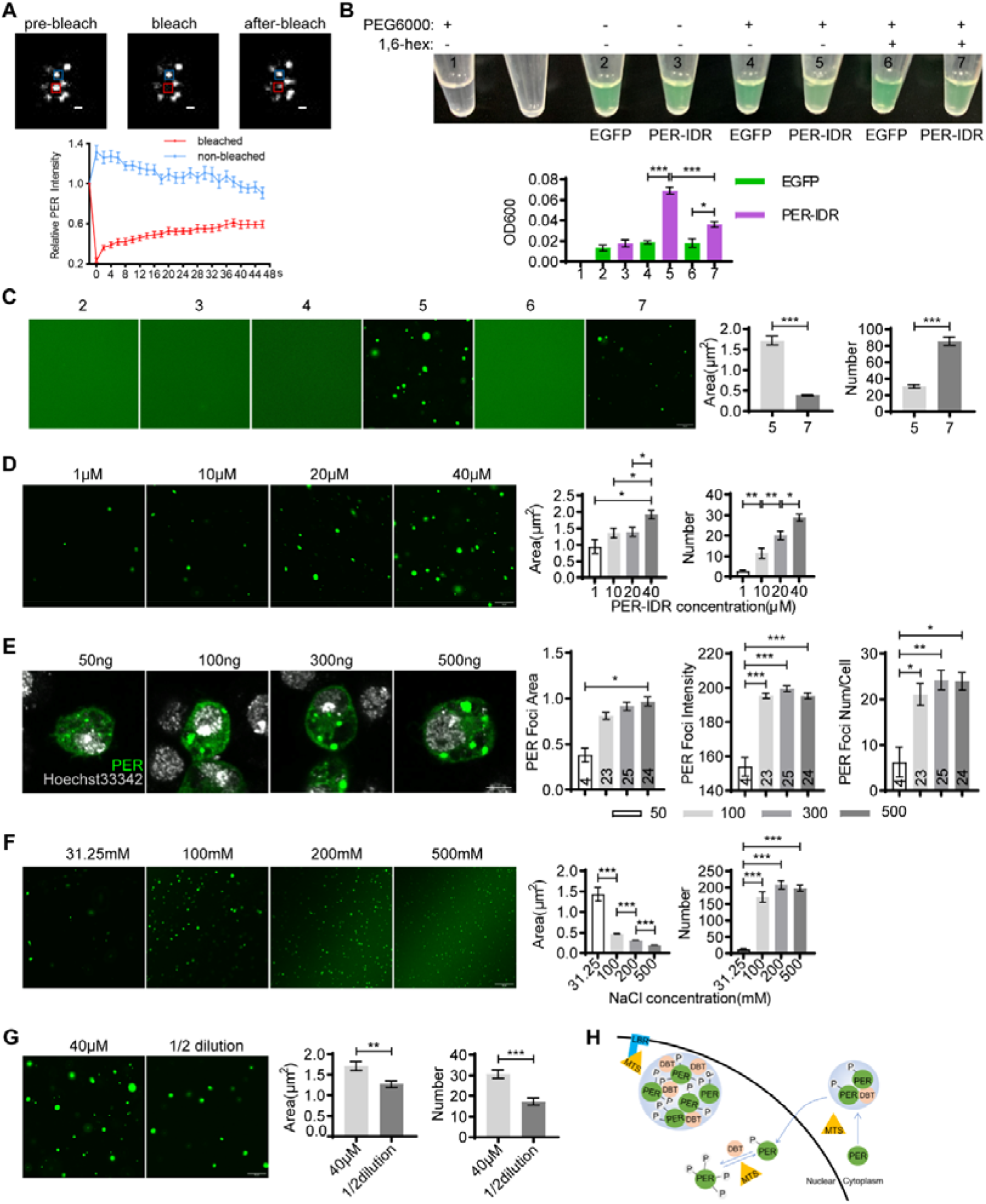
PER foci may be phase-separated condensates. **(A)** Top panel: representative confocal images of the PER foci FRAP assay in S2 cells transfected with pAc-*per*-EGFP for 36hr. Red and blue rectangle represent bleached foci and control, respectively. Scale bar, 1μm. Bottom panel: quantifications of fluorescence intensity. n = 15 cells. **(B)** Top panel: representative image demonstrating the turbidity of 40μM solution of indicated contents with or without 10% PEG6000 and 10% 1,6-hexanediol (simplified as 1,6-hex). Each solution is given a number as an indicator. Bottom panel: plot shows mean optical density 600 (OD600) of the solutions. n = 3. **(C)** Left panel: representative confocal images of protein solutions in (B). Right panel: quantification of the area and number of PER-IDR droplets in the presence PEG6000 or PEG6000 and 1,6-hexanediol. n ≥ 10 fields of view from 3 repeats. Scale bar, 10μm. **(D)** Left panel: representative confocal images of PER-IDR droplets at different protein concentrations. PER-IDR was diluted with buffer to indicated concentrations. Right panel: quantification of the area and number of droplets. n ≥ 10 fields of view from 3 repeats. Scale bar, 10μm. **(E)** Left panel: representative confocal images of S2 cells transfected with indicated dosage of pAc-*per* for 36hr, and immunostained with hoechst33342 (gray) and anti-PER (green). Right panel: quantification of PER foci area, intensity, number per cell. n refers to the number of cells and is indicated on the bars. **(F)** Left panel: representative confocal images of PER-IDR droplets at different NaCl concentrations. Protein concentration used is 10μM. Right panel: quantification of the area and number of droplets. n ≥ 10 fields of view from 3 repeats. Scale bar, 10μm. **(G)** Left panel: representative confocal images of 40μM PER-IDR droplets and 1/2 dilution. Right panel: quantification of the area and number of droplets. n ≥ 10 fields of view from 3 repeats. Scale bar, 10μm. **(H)** Model regarding how LBR, MTS and DBT regulate the accumulation of PER foci. P indicates phosphorylation. Error bars represent SEM. One-way ANOVA, Tukey’s multiple comparison test for (**B, D**-**F**). Student’s t-test for (**C, G**). \**P* <0.05, \*\**P* <0.01, \*\*\**P* <0.001. This figure inclues the following source data: **Figure 4-source data**. The raw data of fluorescence intensity, solution turbidity and area, intensity and number of PER foci/droplets, as well as statistical analyses in ***Figure 4A-G***.

Phase-separated droplets typically scale up in size with increased concentration (Brangwynne, 2013). Here we performed droplet formation assay with varying concentrations of IDR-EGFP ranging from 1μM to 40μM. Indeed, both the size and number of the droplets increase at higher concentrations (Figure 4D). Similarly, the size, intensity and number of PER foci in S2 cells also increase in a dose-dependent manner (Figure 4E).

To investigate the biochemical properties of these IDR-EGFP droplets, we first treated them with 1,6-hexanediol. This substantially reduced the opacity of IDR-EGFP solution, as well as the size and number of the droplets (Figure 4B and C). In addition, we found that 2 hours after the formation of the droplets, they start to exhibit fibrous structure but are still sensitive to 1,6-hexanediol (Figure 4-figure supplement 2C). We next assessed the ability of IDR-EGFP to form droplets under varying salt concentrations which perturb electrostatic interactions. The size of the droplets decreases as salt concentration increases from 31.25mM to 500mN (Figure 4F). These observations implicate that both hydrophobic and electrostatic interactions contribute to the formation of PER IDR condensates.

Lastly, we tested whether the IDR-EGFP droplets are reversible. We first allowed the droplets to form and then the protein concentration was diluted by half. The size and number of the droplets exhibited significant reduction (Figure 4G). These findings demonstrate that PER IDR can form droplets with a distribution of sizes depending on the condition of the system. Once formed, these droplets respond to changes in the system by rapidly altering their size and number. These features are characteristic of phase-separated condensates formed by weak protein-protein interactions (Shin & Brangwynne, 2017).

In summary, these series of results demonstrate that PER IDR can form phase-separated condensates in PEG solution, which may contribute to the formation of PER foci *in vivo*.

## Discussion

LBR is one of the most important proteins in the inner nuclear membrane and is known to play critical roles in tethering heterochromatin to nuclear periphery during development and in cancer cells (Lukasova, Kovar ik, Bac ikova, Falk, & Kozubek, 2017; Nikolakaki et al., 2017; Solovei et al., 2013). Xiao et al reported that knocking down *lbr* in post-mitotic neurons lead to dampening of fly locomotor rhythm, likely due to disturbance of PER foci accumulation (Xiao et al., 2021). Here we found that LBR targets MTS and facilitates its degradation most prominently at CT0, with a similar trend at CT4 and CT8 (Figure 1C and 1D). This corresponds nicely with the peak accumulation of PER foci at CT0 followed by their gradual disappearance (Xiao et al., 2021). Consistently, MTS impedes PER foci accumulation both *in vivo* and in culture cells, which is opposite of the effects of LBR. To our knowledge, LBR has not previously been shown to participate in modulating protein stability. Protein degradation systems have been best described in the cytoplasm and the endoplasmic reticulum, but more recently, the nucleus has also emerged as a key compartment for ubiquitination and proteasome-mediated degradation (Buchberger, Bukau, & Sommer, 2010; Enam, Geffen, Ravid, & Gardner, 2018). Both the nucleus and inner nuclear membrane contain ubiquitination machinery, which could potentially mediate the effects of LBR on MTS (Franic, Zubcic, & Boban, 2021). Further molecular analysis will be required to delineate the detailed mechanism regarding how LBR regulates MTS stability.

The observation that PER form foci-like structure was first observed in the cytoplasm of S2 cells when PER is ectopically expressed (Meyer, 2006). *In vivo*, Xiao et al also found that PER foci first appears in the cytoplasm and then starts to accumulate in the nucleus (Xiao et al., 2021). Our work here strongly suggests that PER foci are phase-separated condensates, and weak interactions of the IDRs of PER promote their formation. These condensates are initially of liquid-like state, but after a while they appear to transition (at least partially) into a “glassy solid” state which are fibrous instead of round (Figure 4-figure supplement 2C). Based on the literature, when condensates are in this state they are more arrested but still reversible (Shin & Brangwynne, 2017). This less dynamic state may also account for the partial fluorescence recovery after photo bleaching the PER foci in S2 cells.

Multiple studies have shown that phosphorylation can regulate phase separation both positively and negatively (Luo, Wu, & Li, 2021). Here we found that although phosphorylation does not appear to be required for PER IDR to form phase-separated condensates *in vitro*, it likely enhances PER foci formation, especially by increasing the size of PER foci. In S2 cells, we observed near perfect co-localization of PER foci and DBT but not MTS, suggesting that DBT may co-exist with PER in phase-separated condensates. Since phosphorylation of PER by DBT is known to play critical roles in determining PER protein stability, we suspect one function of PER foci may be to concentrate PER for more efficient phosphorylation and stability regulation by DBT (J. L. Price et al., 1998). Obviously, much more needs to be done to characterize the contents and function of these foci.

In conclusion, we found PER IDR can form phase-separated condensates, which may contribute to the development of PER foci *in vivo*. Phosphorylation appears to facilitate PER foci accumulation, with DBT increasing their size while MTS reducing their size. LBR promotes PER foci accumulation by destabilizing MTS, adding an extra layer of regulation on PER foci in the nucleus (Figure 4H).

## Materials and methods

### Plasmids

The following plasmids are used in this study: pAc-*per*-V5-His (Lim et al., 2007), pMT-*lbr*-Flag-HA (FMO06243, *Drosophila* Genomics Resource Center), pMT-*mts*-Flag-HA (FMO02385, *Drosophila* Genomics Resource Center), pAc-*per*-EGFP, pMT-mCherry-Flag-His and pMT-*dbt*-mCherry-Flag-His. The open reading freame (ORF) of EGFP was amplified from pEGFP-C2 (#6083-1, Addgene) using upstream and downstream primers located at the BstbI and XbaI cleavage sites, respectively. EGFP ORF was then subcloned into pAc-*per*-V5-His such that EGFP is located at the carboxyl terminal end of *per*. To generate pMT-mCherry-Flag-His, *dbt* with the stop codon was cleaved from pMT-*dbt*-Flag-His (a kind gift from Dr. Joanna Chiu at University of California, Davis) by EcoRI and XbaI, and then mCherry amplified from pCS2+8CmCherry (#34935, Addgene) by PCR was ligated in to the plasmid to obtain pMT-mCherry-Flag-His. The ORF of *dbt* was amplified from pMT-*dbt*-Flag-His and subcloned into pMT-mCherry-Flag-His with the *dbt* ORF at the amino terminal of the mCherry ORF.

### Fly culture

Flies were raised on standard cornmeal food at 25°C and ∼50% humidity under 12h light/12h dark (LD) cycles. The following fly strains were used in this study: *tim*GAL4 (Emery et al., 1998), UAS*dcr2* (V6009, Vienna *Drosophila* Resrouce Center), *w*^*1118*^ (3605, Bloomington *Drosophila* Stock Center), UAS*lbrRNAi* (V110508, Vienna *Drosophila* Resrouce Center), *Pdf*GAL4 (6899, Bloomington *Drosophila* Stock Center), *per*-AID-EGFP (Chen et al., 2018), UAS*mts* (Hannus et al., 2002), UASdn*mts* (Hannus et al., 2002) and UAS*dbt* (J. L. Price, Blau, J., Rothenfluh, A., Abodeely, M., Kloss, B., and Young, M.W., 1998). All fly crosses were carried out at 25°C and male and female offspring were entrained at 25°C for 3 days of LD followed by 1 day of constant darkness (DD) and collected randomly at the indicated time points. For *per*-AID-EGFP experiments only male flies were used.

### Cell culture and transient transfection

*Drosophila* Schneider 2 (S2) cell line was cultured in Schneider’s medium (Gibco) supplemented with 10% fetal bovine serum and 1% penicillin–streptomycin (Thermo Scientific). The cells were plated in a 24-well plate and incubated for 24 hours. siRNA (LBR: CGAAGACAATCTCAAATCT) and negative control were transiently transfected using riboFECTTM CP Regent (RIBOBIO C10511-05) according to the protocol provided by the manufacturer. Plasmids were transiently transfected using Lipofectamine 3000 Transfection Reagent (Thermo Scientific) according to the protocol provided by the manufacturer. Expression of *lbr, mts* and *dbt* under pMT promoter was induced by adding 500 μM CuSO_4_ to the culture upon the completion of transfection and incubation for another 48hr unless specified otherwise. For experiments in which cycloheximide (10 mg/ml; MCE), calyculin A (30nM, MCE) and okadaic acid, sodium salt (5nM, Sigma) were used, they were added at the indicated time post transfection.

### RNA extraction and quantitative real-time PCR (qRT-PCR)

Flies were entrained in LD for 3 days and collected on DD1 at the indicated time points and frozen immediately on dry ice. RNA extraction and qRT-PCR were conducted following our previously published methods (Bu, Chen, Zheng, He, & Zhang, 2020). In brief, total RNA was extracted from fly heads and S2 cells and then subjected to reverse transcription using TransScript One-Step gDNA Removal and cDNA Synthesis SuperMIX (TransGen Biotechnology). All qRT-PCRs were carried out on a Step One Plus Real-Time PCR System (Applied Biosystems, Life Technologies). Primers used are as follows: *rpl32*-f: 5’-TACAGGCCCAAGATCGTGAA-3’, *rpl32*-r: 5’-GCACTCTGTTGTCGATACCC-3’; *mts*-f: 5’-GCGACAAGGCCAAGGAGA3’, *mts*-r: 5’-AGTCGCCCATGAACAGGT-3’. *rpl32* is used as internal control.

### Protein extraction and Western blot

Flies were entrained in LD for 3 days and collected on DD1 at the indicated time points and frozen immediately on dry ice. Fly heads were separated by vortexing and protein extracts were prepared by homogenizing using EB1 (20mM HEPES pH 7.5, 100mM KCl, 5% glycerol, 2.5mM EDTA, 5mM DTT, 0.1% Triton X-100, 25mM NaF) supplemented with complete EDTA-free protease and phosphatase inhibitor cocktail (MCE). S2 cells were harvested and washed once with 1×PBS, and then lysed with EB2 solution (20mM HEPES pH 7.5, 100 mM KCl, 5% glycerol, 5 mM EDTA, 1 mM DTT, 0.1% Triton X-100, 25 mM NaF) supplemented with complete EDTA-free protease and phosphatase inhibitor cocktail (MCE). For cytoplasmic and nuclear extraction, the cytoplasmic fraction was obtained by CER1 (10 mM HEPES pH 7.5, 10 mM KCl, 1.5 mM MgCl2, 0.34 M sucrose, 10% glycerol) supplemented with complete EDTA-free protease and phosphatase inhibitor cocktail (MCE). S2 cells were vortexed vigorously for 15 seconds and incubated on ice for 10 minutes. 3% NP-40 was added to the homogenates and vortexed for 5 seconds. This was followed by incubation on ice for 1 minute and centrifugation at 16, 000 g for 5 minutes at 4°C. The supernatant which is the cytoplasmic fraction was transferred to a new pre-chilled tube. The pellet was suspended with EB2 solution supplemented with complete EDTA-free protease and phosphatase inhibitor cocktail (MCE). The homogenate was vortexed for 15 seconds every 10 minutes for a total of 40 minutes and centrifuged at 16, 000 g for 5 minutes at 4°C. The supernatant acquired is the nuclear fraction.

Equal amounts of protein were run on 10% SDS-PAGE gels and then transferred to nitrocellulose membranes. After incubation with primary antibodies at 4°C overnight, membranes were incubated with secondary antibodies at room temperature for 1 hour. The primary antibodies used were mouse anti-PP2A catalytic subunit (1:1000, Millipore), mouse anti-Hsp70 C7 (1:1000, Abcam), mouse anti-lamin (1:200, Developmental Studies Hybridoma Bank), mouse anti-beta-tubulin, (1:200, Developmental Studies Hybridoma Bank), mouse anti-H2B (1:1000, Abcam). Secondary antibodies used were conjugated either with IRDye 680 or IRDye 800 (LICOR Bioscience) and incubated at a concentration of 1:10000. Blots were visualized with Odyssey Infrared Imaging System (LICOR Biosciences).

### Immunoprecipitation

S2 cells expressing pMT-*lbr*-Flag-HA with or without pAc-*per*-V5-His were harvested 48 hours after transfection. Protein extracts were prepared in EB2 (20 mM HEPES pH 7.5, 100 mM KCl, 5% glycerol, 5 mM EDTA, 1 mM DTT, 0.1% Triton X-100, 25 mM NaF) supplemented with complete EDTA-free protease and phosphatase inhibitor cocktail (MCE) and subsequently incubated with SureBeads to exclude nonspecific binding. Meanwhile, HA antibody (1:50, Cell Signaling Technology) was added to SureBeads and incubated at 4°C for 4 hours. Then, the extracts were incubated with HA antibody-bound beads at 4°C overnight. Beads were magnetized to remove supernatant, washed two times in EB2 and resuspended in 1× loading buffer. Samples were analyzed by Western blotting as described above. 6% SDS-PAGE gel was used for analysis of PER protein. Primary antibodies used were rabbit anti-HA (1:1000; Cell Signaling Technology), mouse anti-PP2A catalytic subunit (1:1000, Millipore) and guinea pig anti-PER (1:1000, a gift from Dr. Joanna Chiu).

### Immunostaining

S2 cells were seeded on cell culture dishes with glass bottom and transfected with indicated plasmids. The cells were washed twice with 1×PBS, fixed with 3.7% paraformaldehyde (PFA) for 10 minutes, and permeabilized with 0.5% Triton X-100 for 15 minutes. Thereafter, the cells were washed with 1×PBS and incubated with blocking buffer (1% BSA, 0.1% Tween-20) for 30 minutes. Subsequently, the cells were incubated with primary antibodies at 4°C overnight. After washing with 0.1% Tween-20 in PBS for four times, the cells were incubated for 1 hour with secondary antibody. Primary antibodies used are as follows: rabbit anti-PER (1:1000, a gift from Dr. Joanna Chiu), mouse anti-HA (1:1000, Medical & Biological Laboratories). Secondary antibodies used were Alexa Fluor 488 goat anti rabbit (1:1000, Abcam) and Alexa Fluor 594 goat anti mouse (1:1000, Abcam). Nuclei were counterstained with hoechst33342 (Beyotime). Finally, the cells were rinsed with 0.1% Tween-20 in PBS four times and mounted with Vectashield Plus Antifade Mounting Medium (Vector Laboratories). The samples were scanned by the Olympus FV3000 confocal microscope with 100×silicone oil objective.

### Live imaging

For fly brain imaging, flies were entrained in LD for 3 days, and dissected on DD1 in chilled Schneider’s medium in less than 10 minutes under low light conditions. A punched double-sided tape was used as a spacer on the slides to prevent flattening of the brains by coverslip. The brains were overlaid with a small amount of Vectashield Plus Antifade Mounting Medium and covered using a coverslip which was sealed with nail polish. The individual z-stack images of s-LNvs were acquired by Olympus FV3000 confocal microscope with 100 × silicone oil objective.

For S2 cell imaging, S2 cells transfected with pAc-*per*-EGFP-V5-His and pMT-mCherry-Flag-His or pMT-*dbt-*mCherry-Flag-His were seeded on cell culture dishes with glass bottom. Hoechst 33342 was added into culture medium 10 minutes before imaging. Live S2 cells were imaged using Olympus FV3000 confocal microscope 100×silicone oil objective with Z series and time-series.

### 1,6-Hexanediol treatment

For treatment with 1,6-hexanediol (MACKLIN), Schneider’s *Drosophila* medium was prepared containing 10% 1,6-hexanediol. Target S2 cells were imaged with Z-stacks and then incubated in 10% 1,6-hexanediol medium for 1 minute, followed by imaging again using the same conditions.

### Fluorescence recovery after photobleaching (FRAP)

FRAP was performed on the Olympus FV3000 microscope with 488 nm laser. PER foci were first identified using a 100×silicone oil objective. Acute light stimulation was achieved by utilizing the 488 nm laser line and stimulation module within the Olympus FV3000 imaging software. Regions of interest (ROIs) to be stimulated were drawn over fields of view prior to image acquisition. Reference ROIs of the same size were drawn adjacent to the cell. Following one baseline image, objects were bleached for 200 milliseconds using 50% laser power (488 nm laser line) and images were collected every 2 seconds post-bleaching. The fluorescence signal measured in the ROIs were normalized to the change in total fluorescence as follows according to a previously published method: I = (T_0_/T_t_) * (I_t_/I_0_) (Phair & Misteli, 2000). T_0_ is the total intensity in the field before bleaching while T_t_ is the total intensity in the field at timepoint t. I_0_ is the average intensity of ROIs before bleaching while I_t_ the average intensity of ROIs at timepoint t.

### Protein purification

cDNAs encoding EGFP and PER-IDR EGFP were cloned into pET30a vector. The sequences were confirmed by sequencing. Plasmids of EGFP and PER-IDR EGFP were transformed into BL21 (DE3) and RosettaTM2 (DE3) E. coli cells respectively. IPTG was added at 0.5mM concentration and incubated for 16 hours at 15°C or 4 hours at 37°C for EGFP and PER-IDR-EGFP, respectively. Pellets from 1L cells culture were collected and sonicated at 15°C at 35% power for 2 seconds at intervals of 4.5 seconds for a total of 1 hour for cells lysis. The lysates were centrifuged at 4°C at 10,000 g for 30 minutes and the supernatants were used to purify target proteins by Ni NTA agarose (GE Healthcare).

### *In vitro* droplet assay

Recombinant protein was added to buffer (50mM Tris pH7.5, 1mM DTT) at varying protein or salt concentrations in the presence or absence of 10% PEG6000 (Sigma) and 10% 1,6-hexanediol. Glass slides were affixed with circular and single-sided tape with a hole drilled in the middle to stop limit liquid from spreading unchecked. The protein solutions were immediately loaded into the hole. The entire process (starting from sample preparation to the completion of imaging) does not exceed 5 minutes to prevent liquid evaporation and droplets from settling on the bottom of the glass.

### Confocal image analyses

Z-stack or time-lapse series were captured by Olympus FV3000 confocal microscope. OIB files were then imported into Fiji as a composite image with a lossless 16-bit resolution per channel. A single Z-plane with the largest foci area or the brightest PER intensity was selected. Each focus was manually annotated for analysis. The mean pixel brightness (arbitrary unit) and geometric area (μm^2^) of the ROI were acquired using built in functions of ImageJ software. For protein droplets assay, the area and number of droplets were analyzed by Fiji automatic counting.

### Statistical analyses

Student’s t-test (Prism Graphpad) was used to analyze the differences between two groups for which data fits normal distribution. One-way ANOVA (Prism Graphpad) was used to compare the differences between multiple groups which have only one explanatory variable. Two-way ANOVA (Prism Graphpad) was used to analyze the difference between two groups with two explanatory variables. Sample size used is consistent with previous literature utilizing similar assays, and samples were selected randomly. All data collected have been included and none was excluded. All replicates used are biological replicates.

## Acknowledgements

We would like to thank Drs. Joanna Chiu, Chunghun Lim, Yong Zhang, Amita Sehgal, and Ravi Allada for kindly providing reagents used in this study. We would also like to thank Min Qin and Dr. Shuguo Sun for helpful advice on phase-separation experiments.

---

**Figure 1-figure supplement 1.**
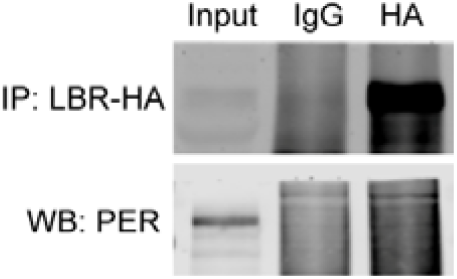
PER does not bind to LBR. Representative Western blots (WB) of protein extracts prepared from S2 cells transfected with pMT-*lbr*-Flag-HA and immunoprecipitates. LBR was immunoprecipitated (IP) with HA antibody, and rabbit IgG was used as control. PER was detected by Western blotting using PER antibody. This figure includes the following source data: **Figure 1-figure supplement 1-source data**. The original Western blots in ***Figure 1-figure supplement 1***.

**Figure 2-figure supplement 1.**
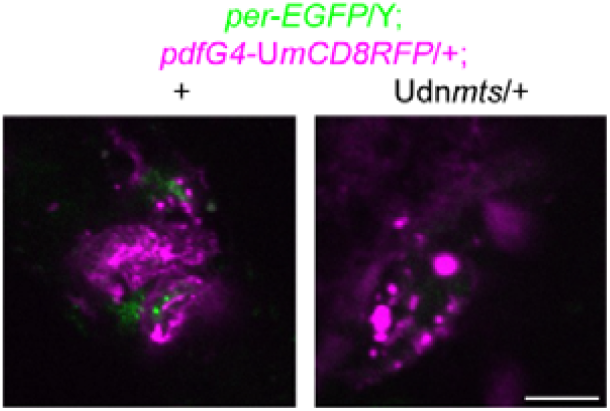
PER foci cannot be observed in dn*mts* expressing flies. Representative confocal images of PER foci in the s-LNvs of *per-*EGFP/Y;*pdf*GAL4*-*UASmCD8RFP*/+* flies over expressing dn*mts* and control flies at CT0 on DD1. Green, PER-EGFP; magenta, mCD8RFP. Scale bar, 5μm. G, GAL. U, UAS.

**Figure 2-figure supplement 2.**
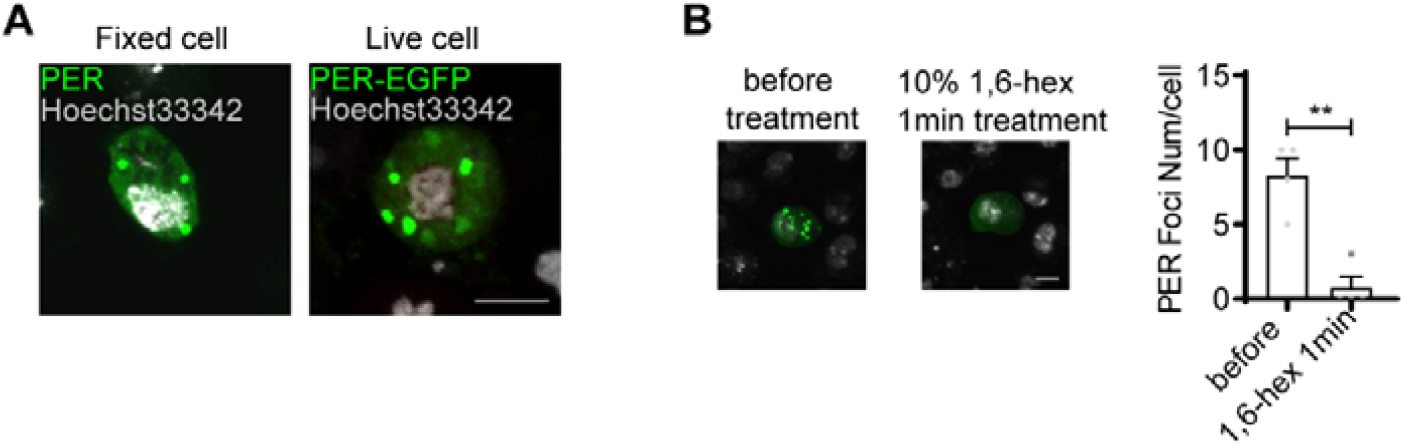
PER foci can be observed in S2 cells. **(A)** Left panel: representative confocal images of PER foci in S2 cells transfected with 100ng pAc-*per* for 36hr and immunostained with hoechst33342 (gray) and PER antibody (green). Right panel: representative confocal images of PER foci in S2 cells transfected with pAc-*per*-EGFP for 36hr and imaged live. **(B)** Representative confocal images of PER foci in S2 cells transfected with 100ng pAc-*per*-EGFP for 36hr and then treated with 10% 1,6-hexanediol for 1min. The plot shows PER foci number per cell. Error bars represent SEM. Student’s t-test. \*\**P* <0.01. Scale bar, 5μm. This figure includes the following source data: **Figure 2-figure supplement 2-source data**. The raw data of PER foci number and statistical analyses in ***Figure 2-figure supplement 2B***.

**Figure 4-figure supplement 1.** FRAP assay of PER foci in PER-EGFP expressing S2 cells.

**Figure 4-figure supplement 2.**
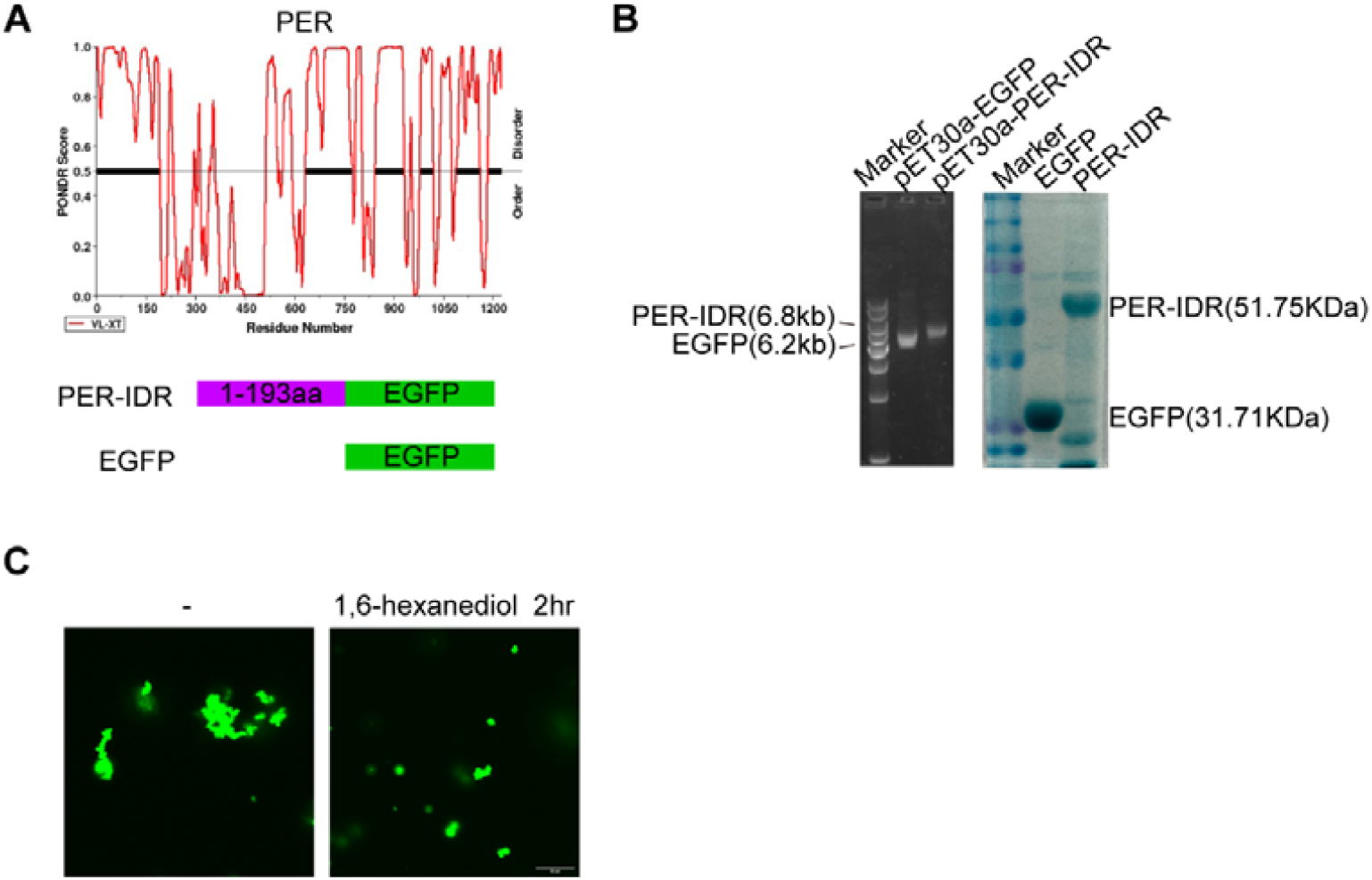
PER-IDR fused with EGFP can form phase-separated condensates in PEG. **(A)** Top panel: disordered region analysis of PER protein using PONDR. Black with bold lines indicate that disordered sequences. Bottom panel: schematics of recombinant protein. **(B)** Electrophoresis gel validation of plasmids encoding the recombinant protein (left) and the purified recombinant protein (right), respectively. **(C)** Representative confocal images of 40μM PER-IDR-EGFP solution treated with 10% PEG6000 in the absence (left) or presence (right) of 10% 1,6-hexanediol treated for 2hr. Scale bar, 10μm.

**Figure 4-figure supplement 3**. Fusion of PER-IDR droplets *in vitro*.

## References

Aguilar-Arnal, L., Hakim, O., Patel, V. R., Baldi, P., Hager, G. L., & Sassone-Corsi, P. (2013). Cycles in spatial and temporal chromosomal organization driven by the circadian clock. Nature structural & molecular biology, 20(10), 1206–1213. doi:10.1038/nsmb.2667

Brangwynne, C. P. (2013). Phase transitions and size scaling of membrane-less organelles. The Journal of cell biology, 203(6), 875–881. doi:10.1083/jcb.201308087

Bu, B., Chen, L., Zheng, L., He, W., & Zhang, L. (2020). Nipped-A regulates the Drosophila circadian clock via histone deubiquitination. EMBO J, 39(1), e101259. doi:10.15252/embj.2018101259

Bu, B., He, W., Song, L., & Zhang, L. (2019). Nuclear Envelope Protein MAN1 Regulates the Drosophila Circadian Clock via Period. Neurosci Bull, 35(6), 969–978. doi:10.1007/s12264-019-00404-6

Buchberger, A., Bukau, B., & Sommer, T. (2010). Protein quality control in the cytosol and the endoplasmic reticulum: brothers in arms. Molecular cell, 40(2), 238–252. doi:10.1016/j.molcel.2010.10.001

Chen, W., Werdann, M., & Zhang, Y. (2018). The auxin-inducible degradation system enables conditional PERIOD protein depletion in the nervous system of Drosophila melanogaster. FEBS J, 285(23), 4378–4393. doi:10.1111/febs.14677

Emery, P., So, W. V., Kaneko, M., Hall, J. C., & Rosbash, M. (1998). CRY, a Drosophila clock and light-regulated cryptochrome, is a major contributor to circadian rhythm resetting and photosensitivity. Cell, 95(5), 669–679. doi:S0092-8674(00)81637-2 [pii]

Enam, C., Geffen, Y., Ravid, T., & Gardner, R. G. (2018). Protein Quality Control Degradation in the Nucleus. Annu Rev Biochem, 87, 725–749. doi:10.1146/annurev-biochem-062917-012730

Franic, D., Zubcic, K., & Boban, M. (2021). Nuclear Ubiquitin-Proteasome Pathways in Proteostasis Maintenance. Biomolecules, 11(1). doi:10.3390/biom11010054

Fu, J., Murphy, K. A., Zhou, M., Li, Y. H., Lam, V. H., Tabuloc, C. A., … Liu, Y. (2016). Codon usage affects the structure and function of the Drosophila circadian clock protein PERIOD. Genes & development, 30(15), 1761–1775. doi:10.1101/gad.281030.116

Grima, B., Chelot, E., Xia, R., & Rouyer, F. (2004). Morning and evening peaks of activity rely on different clock neurons of the Drosophila brain. Nature, 431(7010), 869–873.

Hannus, M., Feiguin, F., Heisenberg, C. P., & Eaton, S. (2002). Planar cell polarization requires Widerborst, a B’ regulatory subunit of protein phosphatase 2A. Development, 129(14), 3493–3503. doi:10.1242/dev.129.14.3493

Li, S., & Zhang, L. (2015). Circadian Control of Global Transcription. Biomed Res Int, 2015, 187809. doi:10.1155/2015/187809

Lim, C., Lee, J., Choi, C., Kim, J., Doh, E., & Choe, J. (2007). Functional role of CREB-binding protein in the circadian clock system of Drosophila melanogaster. Molecular and cellular biology, 27(13), 4876–4890. doi:MCB.02155-06 [pii] 10.1128/MCB.02155-06

Lin, S. T., Zhang, L., Lin, X., Zhang, L. C., Garcia, V. E., Tsai, C. W., … Fu, Y. H. (2014). Nuclear envelope protein MAN1 regulates clock through BMAL1. Elife, 3, e02981. doi:10.7554/eLife.02981

Lukasova, E., Kovar ik, A., Bac ikova, A., Falk, M., & Kozubek, S. (2017). Loss of lamin B receptor is necessary to induce cellular senescence. The Biochemical journal, 474(2), 281–300. doi:10.1042/BCJ20160459

Luo, Y. Y., Wu, J. J., & Li, Y. M. (2021). Regulation of liquid-liquid phase separation with focus on post-translational modifications. Chem Commun (Camb), 57(98), 13275–13287. doi:10.1039/d1cc05266g

Mehsen, H., Boudreau, V., Garrido, D., Bourouh, M., Larouche, M., Maddox, P. S., … Archambault, V. (2018). PP2A-B55 promotes nuclear envelope reformation after mitosis in Drosophila. The Journal of cell biology, 217(12), 4106–4123. doi:10.1083/jcb.201804018

Meyer, P., Saez, L, and Young, M.W. (2006). PER-TIM interactions in living Drosophila cells: An interval timer for the circadian clock Science, 311, 226–229.

Nikolakaki, E., Mylonis, I., & Giannakouros, T. (2017). Lamin B Receptor: Interplay between Structure, Function and Localization. Cells, 6(3). doi:10.3390/cells6030028

Phair, R. D., & Misteli, T. (2000). High mobility of proteins in the mammalian cell nucleus. Nature, 404(6778), 604–609. doi:10.1038/35007077

Price, J. L., Blau, J., Rothenfluh, A., Abodeely, M., Kloss, B., & Young, M. W. (1998). double-time is a novel Drosophila clock gene that regulates PERIOD protein accumulation. Cell, 94(1), 83–95.

Price, J. L., Blau, J., Rothenfluh, A., Abodeely, M., Kloss, B., and Young, M.W. (1998). double-time is a novel Drosophila clock gene that regulates PERIOD protein accumulation. Cell, 94, 83–95.

Renn, S. C., Park, J. H., Rosbash, M., Hall, J. C., & Taghert, P. H. (1999). A pdf neuropeptide gene mutation and ablation of PDF neurons each cause severe abnormalities of behavioral circadian rhythms in Drosophila. Cell, 99(7), 791–802.

Sathyanarayanan, S., Zheng, X., Xiao, R., & Sehgal, A. (2004). Posttranslational regulation of Drosophila PERIOD protein by protein phosphatase 2A. Cell, 116(4), 603–615.

Shin, Y., & Brangwynne, C. P. (2017). Liquid phase condensation in cell physiology and disease. Science, 357(6357). doi:10.1126/science.aaf4382

Solovei, I., Wang, A. S., Thanisch, K., Schmidt, C. S., Krebs, S., Zwerger, M., … Joffe, B. (2013). LBR and lamin A/C sequentially tether peripheral heterochromatin and inversely regulate differentiation. Cell, 152(3), 584–598. doi:10.1016/j.cell.2013.01.009

Stoleru, D., Peng, Y., Agosto, J., & Rosbash, M. (2004). Coupled oscillators control morning and evening locomotor behaviour of Drosophila. Nature, 431(7010), 862–868.

Takai, A., Ohno, Y., Yasumoto, T., & Mieskes, G. (1992). Estimation of the rate constants associated with the inhibitory effect of okadaic acid on type 2A protein phosphatase by time-course analysis. The Biochemical journal, 287 (Pt 1)(Pt 1), 101–106. doi:10.1042/bj2870101

Van Berlo, J. H., Voncken, J. W., Kubben, N., Broers, J. L., Duisters, R., van Leeuwen, R. E., … Pinto, Y. M. (2005). A-type lamins are essential for TGF-beta1 induced PP2A to dephosphorylate transcription factors. Human molecular genetics, 14(19), 2839–2849. doi:10.1093/hmg/ddi316

Xiao, Y., Yuan, Y., Jimenez, M., Soni, N., & Yadlapalli, S. (2021). Clock proteins regulate spatiotemporal organization of clock genes to control circadian rhythms. Proceedings of the National Academy of Sciences of the United States of America, 118(28). doi:10.1073/pnas.2019756118

Zhao, H., Sifakis, E. G., Sumida, N., Millan-Arino, L., Scholz, B. A., Svensson, J. P., … Gondor, A. (2015). PARP1- and CTCF-Mediated Interactions between Active and Repressed Chromatin at the Lamina Promote Oscillating Transcription. Molecular cell, 59(6), 984–997. doi:10.1016/j.molcel.2015.07.019

